# Validating the wearable MUSE headset for EEG spectral analysis and Frontal Alpha Asymmetry

**DOI:** 10.1101/2021.11.02.466989

**Authors:** Cédric Cannard, Helané Wahbeh, Arnaud Delorme

## Abstract

EEG power spectral density (PSD), the individual alpha frequency (IAF) and the frontal alpha asymmetry (FAA) are all EEG spectral measures that have been widely used to evaluate cognitive and attentional processes in experimental and clinical settings, and that can be used for real-world applications (e.g., remote EEG monitoring, brain-computer interfaces, neurofeedback, neuromodulation, etc.). Potential applications remain limited by the high cost, low mobility, and long preparation times associated with high-density EEG recording systems. Low-density wearable systems address these issues and can increase access to larger and diversified samples. The present study tested whether a low-cost, 4-channel wearable EEG system (the MUSE) could be used to quickly measure continuous EEG data, yielding similar frequency components compared to research a grade EEG system (the 64-channel BIOSEMI Active Two). We compare the spectral measures from MUSE EEG data referenced to mastoids to those from BIOSEMI EEG data with two different references for validation. A minimal amount of data was deliberately collected to test the feasibility for real-world applications (EEG setup and data collection being completed in under 5 min). We show that the MUSE can be used to examine power spectral density (PSD) in all frequency bands, the individual alpha frequency (IAF; i.e., peak alpha frequency and alpha center of gravity), and frontal alpha asymmetry. Furthermore, we observed satisfying internal consistency reliability in alpha power and asymmetry measures recorded with the MUSE. Estimating asymmetry on PAF and CoG frequencies did not yield significant advantages relative to the traditional method (whole alpha band). These findings should advance human neurophysiological monitoring using wearable neurotechnologies in large participant samples and increase the feasibility of their implementation in real-world settings.

## I. Introduction

The MUSE (InterAxon Inc.) is a low-cost, off-the-shelf, wearable EEG headset that has two frontal and two temporoparietal (TP) dry active EEG channels. It has been validated for evoked-response potential (ERP) research (i.e., time-domain; [1]) and used in many recent studies [2]–[12]. However, to our knowledge, it has not yet been validated for frequency domain analysis (power spectra on continuous EEG data), with one study showing mixed results [13]. In addition to assessing the validation of MUSE spectral measures, it is relevant to test if the MUSE could be used to estimate clinically- and research-relevant spectral measures, such as the frontal alpha asymmetry (FAA) and the individual alpha frequency (IAF).

Frontal alpha asymmetry (FAA; or frontal EEG asymmetry) refers to the relative difference in log alpha power (8-12 Hz) between the right and the left frontal regions. This spectral measure has been widely used to evaluate participants’ cognitive, emotional, and attentional processes, both as an event-related state response and as a trait during rest [14]–[19]. Because of the inhibitory role of alpha oscillations [20]–[23], relatively greater left than right alpha power is associated with relatively greater right than left cortical activity. In addition, greater activation of the left-frontal cortex relative to the right is related to approach motivation and emotions with positive valence (e.g., happiness, positive urgency), whereas greater activation of the right-frontal cortex relative to the left is associated with the brain processes related to avoidance motivation and negative emotional valence (e.g., depression, anxiety, withdrawal). FAA is suspected to reflect neural processes of the executive control systems and has been source-localized to the frontoparietal network [19].

The individual alpha frequency (IAF) refers to the frequency within the alpha band with dominant spectral power [24]. It is associated with cognitive performance [25], considered a trait-like characteristic of human EEG [26], has high heritability and test-retest reliability [27], [28], and better accounts for interindividual differences in alpha activity [24], [29]. It has been traditionally examined using the peak alpha frequency (PAF) approach, which takes the frequency with the highest alpha power within the alpha band [30]–[32]. However, it has been highlighted that this approach does not perform well in a large portion of the population (up to 44%) that displays absent, ambiguous, or “split” alpha peaks [24], [33]. The alpha center of gravity (CoG) is considered a more robust approach to calculate the IAF by considering the whole alpha power distribution [24].

The IAF may be used to estimate FAA. Since alpha power distribution can fall outside the traditional predefined range (8-13 Hz) for some individuals [32], asymmetry scores based on the IAF (instead of the traditional band) might better address interindividual differences and might therefore provide more accurate asymmetry indexes method for research and clinical applications [34], [35].

IAFs and FAA seem like promising candidate measures for wearable EEG systems, as they require simple calculations in the frequency domain and a few EEG channels covering the frontal regions of each hemisphere. While these measures have not been validated using these systems against research-grade EEG, wearable EEG systems have been used extensively over the past few years to measure frontal asymmetry, suggesting this measure is well-suited for these technologies [2], [36]–[45]. Wearable systems, when reliable, can offer advantages for researchers through easeful EEG data collection for over large samples, increased access to populations that are hard to study with conventional systems (e.g., children, elderly, patients), reduced hardware and software costs, and facilitated EEG research in real-world environments by increasing subjects’ mobility and streaming the data wirelessly [46].

However, there is still a lack of validation of the data collected by such devices and the interpretation of the results based on the literature based on conventional higher-density systems and different referencing methods (i.e., linked-mastoids or average reference). The reference method implemented for low-density wearable systems is of particular importance when considering measuring EEG asymmetry [47], [48]. Both IAF and FAA are promising EEG measures for neurofeedback applications [35], [49], which would benefit from mobile data collection.

The present study tested whether the 4-channel wearable MUSE EEG system can quickly measure continuous EEG data with a maximum of 5-minute set-up and data collection time, that would yield quantifiable frequency components comparable to research-grade systems and if it can extract clinically relevant measures such as IAF and FAA.

## II. METHODS

### A. Participants

Participants for this study were 40 English-speaking adults in the San Francisco Bay area. Exclusion criteria were: aged younger than 18 years old, unable to read, having an acute or chronic illness that interfered with the completion of the experiment, or being unable to sit on a chair for about 30 minutes. Participants had their EEG recorded with a 64-channel EEG system at the laboratory for another study (~2h session) and were asked if they wanted to volunteer a few more minutes of their time for an additional ~5 minutes EEG recording using the wearable headset. They were compensated only for their participation in the initial study. They gave informed consent, and the study was approved by the IONS Institutional Review Board.

### B. EEG data collection procedures

EEG data were collected with the active dry MUSE 1 (version 2016) at 256 Hz and a 64-channel gel-based BIOSEMI Active 2 system (BIOSEMI Inc.) at 512 Hz. Simultaneous recording of both systems was not possible due to their configurations. The MUSE data were recorded first, and then the BIOSEMI data about 30 minutes later, which corresponded to the time necessary to set up the BIOSEMI equipment and optimize channel impedance). A comparison of the two systems’ hardware specifications can be found in Table 1. For both systems, the participants’ skin was cleaned with alcohol wipes at electrode sites before positioning the headband/head cap.

**Table I.** HARDWARE SPECIFICATIONS OF EACH SYSTEM

|  | Biosemi Active Two | InteraXon MUSE |
| --- | --- | --- |
| **Electrode montage** | 64 wet active electrodes (10-20 system) | 4 dry active electrodes (AF7, AF8, TP9, TP10)<sup>b</sup> |
| **Sampling rate** | 512 Hz | 256 Hz |
| **Resolution** | 24 bits | 12 bits |
| **Active electrode system** | Passive DRL and active CMS located around POz | Passive DRL and active CMS located at Fpz |
| **Head sizes** | 3 different head cap sizes covering 54-62 cm | Adjustable headband, 52-60 cm range |
| **Recording apparatus** | Optic fiber and amplifier, MacBook Pro, Actiview Software | Bluetooth on a low-cost Chromebook, Mind Monitor App |
| **Reference** | Reference free<sup>a</sup> | Fpz |

<sup>a</sup> Data is reference-free at data collection time. A reference must be chosen when importing the data. Not choosing a reference led to a 40 dB loss of signal-to-noise (SNR) ratio.
<sup>b</sup> Approximate positions.

#### MUSE

A thin layer of water was applied to the dry electrodes with a sponge for both the frontal metallic sensor and the conductive silicone rubber mastoid sensors behind the ears to decrease the impedance and increase signal quality. The MindMonitor App [50] running on a Chromebook laptop was used to record the EEG signal and check electrode contact (a colored circle for each electrode was filled when the software deemed the connection acceptable). Visual examination of the raw EEG waveforms was also performed while participants were asked to generate eye blinks to provide an additional index of signal quality. The headset position was adjusted if the signal was judged too noisy by visual inspection of the data.

#### BIOSEMI

Highly conductive electrolytes SignaGel was injected into the electrode sites of the BIOSEMI head cap. BIOSEMI active electrode offsets were kept below offset 20 using the Actiview software.

#### MUSE and BIOSEMI

Recordings were performed at the same location within the recording room, minimizing the differences in terms of potential electrical artifacts from the environment. One minute of data was recorded with eyes open gazing at the computer screen in front of them, and one minute was recorded with eyes closed. Half the participants did eyes open before eyes closed, and the other half did the reversed order to avoid carry-over effects. Participants were instructed to sit still on a chair, limit their movements, and focus their attention on their breath by counting each inhalation/exhalation cycle. In this manuscript, we only process eyes’ closed data.

### C. EEG data preprocessing

BIOSEMI data were imported into the EEGLAB processing software (v2021.1; [51]) using the BIOSIG plugin (v3.7.5). MUSE data were imported using the MUSEMonitor (v3.2) plugin of EEGLAB. BIOSEMI data were downsampled to 256 Hz. Raw data were high-pass filtered with EEGLAB’s linear non-causal Finite Impulse Response (FIR) filter of the FIRFILT (v2.4) plugin (filter order = 1129; transition bandwidth = 0.75 Hz; passband edge = 0.75 Hz; −6 dB cutoff frequency = 0.375 Hz). No low-pass filter was used.

Files were inspected visually for abnormal channels (bad connection, impedance, very high noise, flat sections from disconnections, etc.) and artifactual segments (eye and muscle artifacts, high-frequency bursts, etc.). Artifactual regions and channels were manually rejected. MUSE data files with at least 1 visually abnormal channel were removed. If the BIOSEMI or the MUSE file was shorter than 45 s, the participant data was also excluded from further analysis. Using these criteria, three out of 40 data files were excluded.

The traditional method to compute frontal alpha asymmetry (FAA) is to calculate the difference in log-transformed alpha power between the frontal electrodes F7 and F8 on 64-channel EEG data [47], [48]. While the linked-mastoids reference method has been used extensively in the EEG asymmetry literature, average-referencing was shown to be preferable to estimate FAA [47]. Thus, spectral measures were obtained on BIOSEMI data referenced to the average (called the “*average-ref montage*” in this study) and on BIOSEMI re-referenced to mastoids (called the *“mastoid-ref montage”*). With 4 electrodes, an average reference is not meaningful for the MUSE system since it requires a whole-head electrode coverage. The default reference channel for the MUSE is Fpz which is close to the frontal channels AF7 and AF8, and leads to low signal amplitude on these channels. Thus, the MUSE frontal channels were re-referenced to the TP9/TP10 mastoid electrodes (the two other channels available on the MUSE), termed in this study the “*mastoid-ref montage*” (AF7 and AF8 with linked mastoid reference)*”*. This reference method has been widely used in the asymmetry literature (e.g., [47], [52]).

To assess if spectral measures obtained with the MUSE *mastoid-ref montage* are reliable and interpretable in terms of underlying neural activity, we tested whether they were comparable to those obtained with the BIOSEMI *mastoid-ref montage* and the BIOSEMI *average-ref montage.*

### D. Power spectral density (PSD)

Power spectral density (PSD) was computed using the *pwelch* function in MATLAB 2021a (The MathWorks Inc., MA, United States) for each EEG channel on 4-second hamming windows, with 50% overlap and 200% padding (taking into account data discontinuity due to excluded artifactual regions). The mean was removed from PSD data, and they were converted to decibels (10*Log_10_(power)) [48]. Mean PSD was extracted for each frontal channel for each frequency band: delta (1-3 Hz), theta (3-7 Hz), alpha (8-13 Hz), beta (14-30 Hz), and gamma (>30 Hz). Then, the average between the two channels was used for analyses.

### E. Individual alpha frequency (IAF)

Both the peak alpha frequency (PAF) and the alpha center of gravity (CoG) were estimated using the open-source and automated *restingIAF* toolbox (v1.0.2; [24]). This method uses curve-fitting algorithms, zero-crossing, and Savitzky-Golay Filter (SGF) smoothing techniques (same parameters as above for PSD estimation, a minimum of 1 required channel to estimate PAF and CoG, and the default values for the other parameters).

### F. Frontal alpha asymmetry

Three methods were used to calculate alpha asymmetry:

- *Traditional method:* the difference between the frontal channels on alpha power (in dB) averaged over the 8-13 Hz band (*mean_alpha_right - mean_alpha_left*).
- *PAF-asymmetry.* Same as above but on power at the peak alpha frequency (PAF).
- *CoG-asymmetry*. Same as above but on power at the alpha center of gravity (CoG).

### G. Internal consistency reliability

Previous research showed that reliable asymmetry values can be obtained with as little as 80 seconds of data [53]. To confirm internal consistency reliability of the asymmetry measures with the different montage methods and with very short segments of data (45 seconds for the shortest file after data cleaning), mean alpha power and FAA (traditional method only) were also computed for each montage on eleven 4-s blocks of data (mean for each block). Internal consistency reliability of alpha PSD and FAA was evaluated using Cronbach’s standardized alpha on the blocks of spectral data [54], [55]. Values >.8 indicate high internal consistency reliability and <.3 indicate low internal consistency reliability; [53].

### H. Statistics

All spectral measures were compared using the skipped Pearson correlation from the open-source Robust Correlation MATLAB toolbox [56]. Skipped Pearson correlations detect and remove bivariate outliers using the minimum covariance determinant (MCD) estimator, and better control for the type I error by accounting for their deletion when testing for significance, and by using bootstrapped *95% confidence intervals* (*CI*; [56]–[58]). If the *CI* encompasses 0, then the null hypothesis (*H0*) of independence cannot be rejected. This approach is less sensitive to heteroscedasticity (i.e., change in the spread of the residuals over the range of measured values leading to biased results) and therefore more robust against the type I error [56], [57]. Rejections of *H0* at the 95% confidence level (i.e., significant correlations) are reported next to the skipped Pearson correlation *r coefficient* scores with *\** (i.e., *p < 0.05*). Bivariate outliers correspond to the red observations in the plots. The red line correspond to the least square fit line, and the red shaded areas correspond to the *95% CI*.

## III. RESULTS

### A. Internal consistency reliability

The following Cronbach’s alpha scores were obtained for frontal alpha power (.98 - BIOSEMI *average-ref montage;* .95 - MUSE *mastoid-ref montage)* and frontal alpha asymmetry (.67 - BIOSEMI *average-ref montage;* .76 - MUSE *mastoid-ref montage).*

### B. Power spectral density (PSD) for each frequency band

The averaged PSD of each frequency band was first compared between the BIOSEMI *mastoid-ref montage* and the MUSE *mastoid-ref montage*. All frequency bands were significantly correlated between the two montages: delta (1-3 Hz, r = .59*, CI [0.38, 0.75]), theta (3-7 Hz, r = .73*, CI [0.55, 0.85]), alpha (8-13 Hz, r = .87*, CI [0.77, 0.93]), beta (14-30 Hz, r = .84*, CI [0.70, 0.91]), and gamma (>30 Hz, r = 0.48*, CI [0.19, 0.69]). These results are plotted in Fig. 1.

**Fig. 1.**
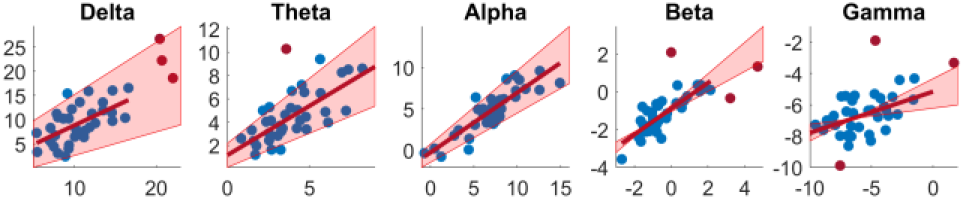
Correlations between BIOSEMI (mastoid-ref montage) and MUSE (mastoid-ref montage) of mean power spectral density (PSD) for each frequency band: delta (1-3 Hz), theta (3-7 Hz), alpha (8-13 Hz), beta (14-30 Hz), and gamma (> 30 Hz). All frequency bands were significantly correlated. Statistics are reported in the text of the Results section. Red dots are bivariate outliers accounted for by the skipped Pearson correlations. The red line is the least-squares fit line. Shaded areas are the 95% confidence intervals. The power spectral density (PSD) unit is deciBels (10*log10(μV^2^/Hz)).

Correlations between PSD estimates from MUSE *mastoid-ref montage* and those from BIOSEMI *average-ref montage* are reported in Fig. 2. Significant correlations were observed for the delta (r = .47*, CI [0.19, 0.69]), the theta (r = .63*, CI [0.43, 0.78]), the alpha (r = .80*, CI [0.65, 0.90], and the beta (r = .74*, CI [0.58, 0.86]) bands. However, the correlation was not significant for the gamma band (r = .17, CI [-0.13, 0.50]).

**Fig. 2.**
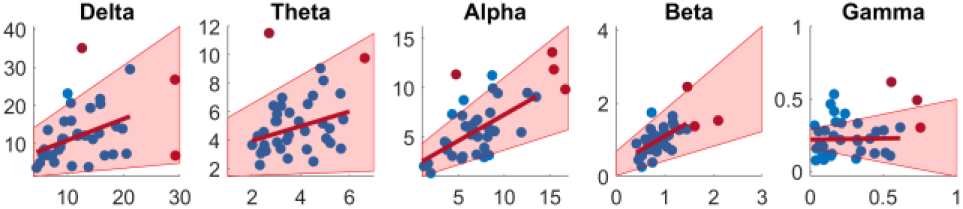
Correlations between BIOSEMI (average-ref montage) and MUSE (mastoid-ref montage) of mean power spectral density (PSD) for each frequency band: delta (1-3 Hz), theta (3-7 Hz), alpha (8-13 Hz), beta (14-30 Hz), and gamma (> 30 Hz). All frequency bands except gamma were significantly correlated. Statistics are reported in the text of the Results section. Red dots are bivariate outliers accounted for by the skipped Pearson correlations. The red line is the least-squares fit line. Shaded areas are the 95% confidence intervals. The power spectral density (PSD) unit is deciBels (10*log10(μV^2^/Hz)).

### C. Individual alpha frequency (IAF)

IAFs estimated on BIOSEMI *mastoid-ref montage* were significantly correlated with those obtained on MUSE *mastoid-ref montage* (Fig. 3, left), for both PAF (r = .91*, CI [0.79, 0.97]) and CoG (r = .78*, CI [0.64, 0.88]). However, PAF could not be estimated on BIOSEMI for 7 files, and for MUSE on 13 files. CoG could not be estimated on BIOSEMI data for 5 files and on MUSE data for 4 files.

**Fig. 3.**
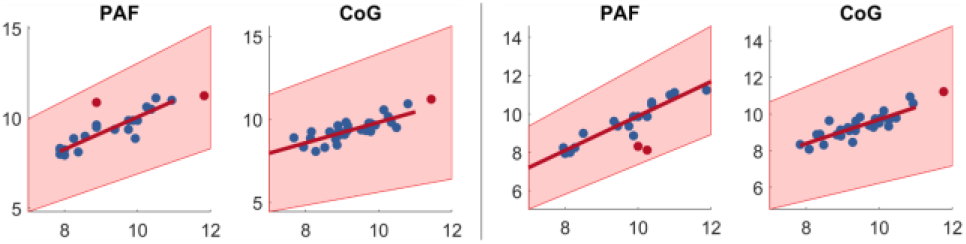
Left: Correlations between BIOSEMI (mastoid-ref montage) and MUSE (mastoid-ref montage) of individual alpha frequency (IAF). Right:between BIOSEMI (average-ref montage) and MUSE (mastoid-ref montage). All estimates using both the peak alpha frequency (PAF) and the alpha center of gravity (CoG) were significantly correlated between the two systems and montages. Statistics are reported in the text of the Results section. Red dots are bivariate outliers accounted for by the skipped Pearson correlations. The red line is the least-squares fit line. Shaded areas are the 95% confidence intervals. The power spectral density (PSD) unit is deciBels (10*log10(μV^2^/Hz)).

Correlations between IAF for the BIOSEMI *average-ref montage* and the MUSE *mastoid-ref montage* (Fig. 3, right) were also significant for both estimation methods: PAF (r = .95*, CI [0.86, 0.98]) and CoG (r = .84*, CI [0.69, 0.93]).

However, the automated algorithms could not detect the PAF for 18 files (11 on BIOSEMI data and 13 on MUSE data) and the CoG for 6 files (5 for BIOSEMI and 4 for MUSE).

### D. Frontal alpha asymmetry (FAA)

The three methods to compute FAA were significantly correlated between BIOSEMI and MUSE with the same mastoid-ref montage: traditional asymmetry (r = .67*, CI [0.40, 0.93]), PAF-asymmetry (r = .35*, CI [0.7, 0.62], CoG-asymmetry (r = 0.42*, CI [0.05, 0.69]). These results are plotted in Fig. 4.

**Fig. 4.**
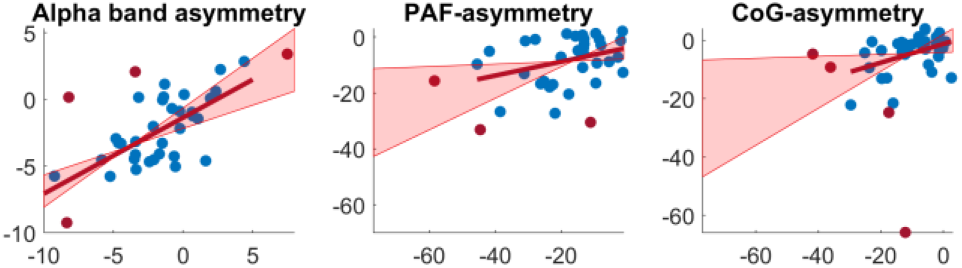
Comparison of frontal alpha asymmetry measures from BIOSEMI mastoid-ref montage and MUSE mastoid-ref montage. The three forms of frontal alpha asymmetry were significantly correlated between the two systems. Statistics are reported in the text. Red dots are bivariate outliers accounted for by the skipped Pearson correlations. The red line is the least-squares fit line. Shaded areas are the 95% confidence intervals. The power spectral density (PSD) unit is deciBels (10*log10(μV^2^/Hz)).

Finally, FAA measures were compared between the BIOSEMI *average-ref montage* and the MUSE *mastoid-ref montage* and are plotted in Fig. 5. FAA calculated on the average power over the whole alpha band (i.e., traditional method) was significantly correlated (r = .37*, CI [0.06, 0.60]). However, asymmetry scores calculated on power at the PAF (r = .12, CI [-0.24, 0.44]) and at the CoG (r = .26, CI [-0.02, 0.55]) were not significantly correlated.

**Fig. 5.**
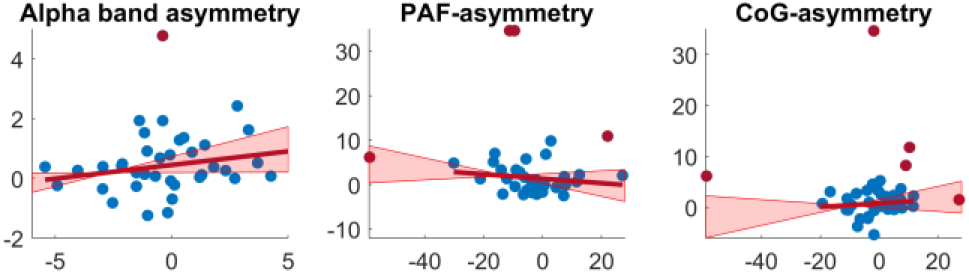
Comparison of frontal alpha asymmetry measures from BIOSEMI average-ref montage and MUSE mastoid-ref montage. Alpha asymmetry calculated using the traditional method (on average power over the whole alpha band) was significantly correlated between the two systems. However, asymmetry scores calculated on the PAF and CoG power were not significantly correlated. Statistics are reported in the text. Red dots are bivariate outliers accounted for by the skipped Pearson correlations. The red line is the least-squares fit line. Shaded areas are the 95% confidence intervals. The power spectral density (PSD) unit is deciBels (10*log10(μV^2^/Hz)).

## IV. Discussion

### A. Results and interpretations

When comparing MUSE *mastoid-ref montage* with BIOSEMI *mastoid-ref montage*, all spectral measures significantly correlated, indicating that this low-cost wearable EEG system can accurately capture these frequency components and that interpretations are in line with the literature using *mastoid-ref montages* can be made. However, correlation coefficients and CIs indicate that the traditional method to calculate frontal alpha asymmetry should be preferred relative to the PAF- and CoG-asymmetry methods.

When comparing MUSE *mastoid-ref* montage with BIOSEMI *average-ref montage*, PSD (in all frequencies beow 30 Hz), IAF, and FAA (traditional method) were significantly correlated, indicating that the MUSE can be used to examine these measures and interpret the findings in line with the literature using the *average-ref montages* (i.e., F7 and F8 sites referenced to average). PAF- and CoG-asymmetry measures were not significantly correlated, indicating they should only be interpreted in the *mastoid-ref montage* context.

These latter findings may suggest that:

1. The automated toolbox used for IAF-estimation does not perform well on low-density sparse montages and is better suited for higher density montages (since it can use neighboring channels to improve detection performance; [24]). Channels referenced to average may have contained alpha spectral components from other channels that were not captured by the *mastoid-ref montage*. IAF measures (PAF and CoG) could not be estimated for some files, which could have reduced statistical power compared to the traditional measures. However, the superior performance of the CoG method compared to the PAF method was apparent since it was able to find IAF in many more participants.
2. The traditional asymmetry method is more robust and grounded in theory (independently of the montage). Previous research suggested that EEG asymmetry is influenced by different neural processes between the lower and the upper frequencies of the alpha band [32]. Thus, while IAFs better account for interindividual differences and are associated with some cognitive processes (e.g., memory), they might reflect different underlying neural processes than those underlying alpha asymmetry (e.g., executive control, attention, emotion regulation). Thus IAF-asymmetries might not be well-suited for asymmetry calculation.

### B. Limitations

The first limitation of this study is the 30 minutes difference between the two recordings. Mental states may likely have changed between the two recordings. However, correlations were still significant when comparing the MUSE and the BIOSEMI with the same *mastoid-ref montage*, suggesting trait spectral components were still captured. Ideally, both types of data should have been recorded simultaneously using markers to synchronize the data at the millisecond resolution. While this was not possible for this study, future studies should aim to record both systems simultaneously.

Second, FAA during rest was previously estimated to vary ~60% from trait influence and 40% from state influences [59], the former being the target measure in this study. While internal consistency reliability of asymmetry measures was relatively high, more variation and lower values were observed compared to the internal consistency reliability of the alpha power data (as in previous publications; [53]). Increasing the data length (e.g., 3 minutes of artifact-free data) might increase the trait influence by reducing the fluctuations due to state influences, and in turn, increase internal consistency reliability. We purposely used short segments to determine if they could be easily and reliably used in experimental and clinical conditions, but we did not compare different data lengths and their impact on these trait EEG measures. Future studies should compare asymmetry measures from a clinical system and a low-cost wearable system (as in this study) with longer data lengths to address this potential limitation.

The absence of correlation in the higher frequencies (PSD > 30 Hz) when comparing MUSE with BIOSEMI *average-ref montage* but not *mastoid-ref montage* may suggest that these frequencies may reflect field potentials from other brain processes when referenced to average than those captured with the *mastoid-ref montage*. Thus, these frequencies should only be interpreted in the *mastoid-ref montage* context when uing this system.

### C. Recommendations for research and clinical MUSE recordings

Recommendations for using the MUSE in future clinical and experimental research are as follows:

- Eye’s closed recordings of at least 1 minute (ideally 5 if time allows), corresponding to a total preparation and recording time of about 3 minutes (about 8 for 5-minute recordings).
- Cleaning the participants’ skin with alcohol wipes and wetting the dry electrodes to reduce impedance.
- Manual exclusion of channels and bad data portions after data collection (or validation of an automated method on this system’s signal).
- Re-referencing the data to linked mastoid electrodes (i.e., TP9/TP10).
- Using measures found to be reliable with this system: PSD<30 Hz, traditional FAA, and the IAF (in particular the CoG).
- Robust statistical methods should be used to better account for outliers and higher noise in the spectral estimates that might occur more frequently using this montage and system (e.g., skipped correlations, iteratively reweighted least squares regressions or weighted least squares regressions).

## V. Conclusion

Our study validates the use of the low-cost MUSE headset for accurately and reliably measuring PSD, IAFs, and FAA (calculated on the whole band). This system can help advance human neurophysiological monitoring techniques on large datasets using wearable neurotechnologies and increase the feasibility of their implementation into real-world applications.

